# Fast single molecule localization microscopy in 3D

**DOI:** 10.64898/2026.09.29.755394

**Authors:** Edward S. Allgeyer, Jennifer H. Richens, Jeanne Lefévère-Laoide, Ewa K. Paluch, George Sirinakis, Daniel St Johnston

**Affiliations:** The Gurdon Institute & the Department of Genetics, University of Cambridge, Tennis Court Road, Cambridge, CB2 1QN, UK; Department of Physiology, Development and Neuroscience, University of Cambridge, Downing Street, Cambridge, CB2 3DY, UK

**Author notes:** Corresponding Authors: George Sirinakis, Daniel St Johnston. These authors contributed equally to this work (Edward S. Allgeyer, George Sirinakis).

## Abstract

Super-resolution methods, such as single molecule localisation microscopy (SMLM)^1–3^, can determine the distribution of specific proteins at the nanometre scale. However, three-dimensional (3D) SMLM is typically limited to thin sections and requires prohibitively long acquisition times to image complete cells. Here we introduce a 3D SMLM imaging modality, Intrinsic3D, that combines the speed and optical sectioning of oblique plane microscopy (OPM)^4,5^ with the nanometre resolution of SMLM to determine the position of single molecule emitters throughout complete volumes more than an order-of-magnitude faster over regular DNA-PAINT. By scanning the oblique light sheet across the volume and combining localization information from two or more adjacent planes, an emitter’s x, y and z position is determined, eliminating the need for post-processing steps to stitch multiple image stacks together. We demonstrate resolution and speed by imaging nuclear pore complexes in U2OS cells and *Drosophila* egg chambers. As a proof-of-concept, we map the 3D distribution of the cell-adhesion protein *D*E-cadherin in the *Drosophila* follicular epithelium and show that spot adherens junctions are enriched near tricellular junctions and contain more E-cadherin than the apical adherens junctions, suggesting that spot junctions play a wider role in cell-cell adhesion than previously appreciated. Finally, we examine the organization of Gliotactin, a tricellular septate junction protein relative to E-cadherin using exchange-PAINT. These investigations demonstrate that Intrinsic3D delivers rapid 3D protein mapping in whole cells. Combining Intrinsic3D with existing multiplex and resolution enhancing methods (e.g. exchange-PAINT and RESI) will enable multicolour, 3D *in situ* spatial proteomics in tissue samples and bridge the gap between SMLM and systems-level cell biology.

## Main

Single molecule localization microscopy^1–3^ (SMLM) provides specific protein distribution information with nanometer resolution and is poised to become critical for the emerging field of spatial proteomics. Early two-dimensional (2D) SMLM approaches quickly evolved into three-dimensional (3D) implementations through pioneering developments including point spread function (PSF) engineering (e.g. astigmatism^6^, double helix PSF^7^, tetrapod PSF^8^), light field microscopy^9^, bi-plane detection^10^, and 4Pi interferometry^11–13^. These approaches typically encode the axial position of the single-molecule emitters in 2D images.

A prerequisite for SMLM is ‘blinking’ which may be produced by photophysical switching or short transient binding in the case of DNA-PAINT^14^, where a freely-diffusing probe conjugated to a short oligonucleotide (imager strand) transiently binds to a complementary oligonucleotide (docking strand) attached to a protein of interest. When bound, the imager produces an effective blink that can be localized. DNA-PAINT offers predictable binding/blinking kinetics for image quantification, the ability to massively multiplex, and mitigates photobleaching. However, freely diffusing imager probes in solution create a large fluorescent background compromising blink detection and localization, especially when imaging deeper in cells and tissues where subtle PSF shape changes become difficult to detect. As such, DNA-PAINT has largely been limited to applications close to the cover glass. To overcome this, imaging modalities that provide optical sectioning, such as line scanning or spinning disk confocal have been employed^15,16^. However, these modalities only use a fraction of the available exposure time (duty cycle), reducing the number of collected photons^17^. Light sheet microscopy offers an attractive alternative by providing optical sectioning while maintaining high photon collection efficiency. Combining lattice light sheet, with an astigmatic PSF, Legant *et al.* were able to reconstruct 3D volumes of cells but required acquisition times ranging from 3-10 days^18^. Oblique plane microscopy^4,5^ (OPM), a variant of light-sheet microscopy that uses a single high NA objective lens and is compatible with conventional sample mounting methods (slides and dishes), has also been successfully adapted for SMLM imaging^4,19,20^. Crucially, this method enables fast volumetric imaging without mechanical agitation, as the light-sheet can be scanned across the sample using galvanometric mirrors^21^.

Here we combine the unique features of OPM with localization microscopy to introduce Intrinsic3D, a method for SMLM imaging that intrinsically determines the position of single molecule emitters throughout a complete volume with more than an order-of-magnitude increase in acquisition speed. Instrinsic3D discretely samples a single molecule’s PSF at consecutive and specific locations during a single blinking event to: (1) laterally (x, y) localize the molecule using established methods, and (2) measure the number of photons at each sampling point to determine the axial (z) position. Because the PSF sampling is performed at predetermined locations throughout the sample volume on the oblique plane, which intrinsically includes depth information, emitters are localized in all three-dimensions (x, y, z) throughout the whole imaging volume, eliminating the need for stitching multiple image stacks in post processing. Unlike previous 3D methods, the axial (z) position determination does not require the detection of subtle PSF shape changes or photon partitioning schemes to infer axial position. Thus, it eliminates complexity and maintains good performance over greater imaging depths in cells and tissues with low photon numbers.

### Using an oblique light-sheet to intrinsically localize molecules in 3D

The implementation of Intrinsic3D is illustrated in **Fig. 1** where an oblique light-sheet illuminates a plane along the Y and S axes at an angle of ∼26° with respect to the X axis (**Fig. 1a**). By stepping the oblique light-sheet along the X direction, multiple sequential images of an emitter’s position on the YS plane are captured. Each step probes the PSF at a different z-position (**Fig. 1b**). Localizations from each YS image are combined to determine the (x, y) position as a localization precision weighted average while the number of photons vs. light-sheet position is fit with a Gaussian model to extract the emitter’s z-position (**Methods Section** and **Fig. 1c**). The oblique light-sheet geometry and its extent along the S and Y axes enables the localization of emitters through complete volumes in a single scan without the need for stitching and/or de-skewing of large image stacks in post processing (**Fig. 1d**). Given that Intrinsic3D relies on stepping a light-sheet to sample an emitter’s PSF multiple times during a single blinking event, the camera exposure, scan length and scan step size must be jointly optimized. For example, shorter camera exposure times expedite imaging but collect fewer photons and reduce the viable measurement range around an emitter, ultimately degrading the localization precision (**Supplementary Fig. 1a-c**). To better understand these trade-offs, we simulated an Intrinsic3D experiment. Simulations showed that increasing the measurement range around an emitter’s position drastically increased z localization precision (**Supplementary Fig. 1c**), while increasing the number of consecutive measurements yields a further, but more modest, improvement in localization precision along all three dimensions (**Supplementary Fig. 1c**). We experimentally confirmed these results by imaging DNA origami (**Fig. 1e-g, Supplementary Fig. 2**). Furthermore, simulations showed that to maximize the number of localized binding events in the imaging volume, large step sizes were favourable (**Supplementary Fig. 1d-i**). The optimal imaging parameters ultimately depend on the application-specific balance between acquisition speed, localization precision and sample characteristics. Here, with an average probe binding time of ∼0.2-1 s, a camera exposure time of 20 ms ensured a minimum of two PSF measurements per binding event with an X step size of 700 nm (**Supplementary Fig. 1f-i**). These conditions were used for biological imaging as it minimized acquisition time without compromising localization precision compared to previous 2D measurements on similar tissue specimens^16,22,23^.

**Figure 1.**
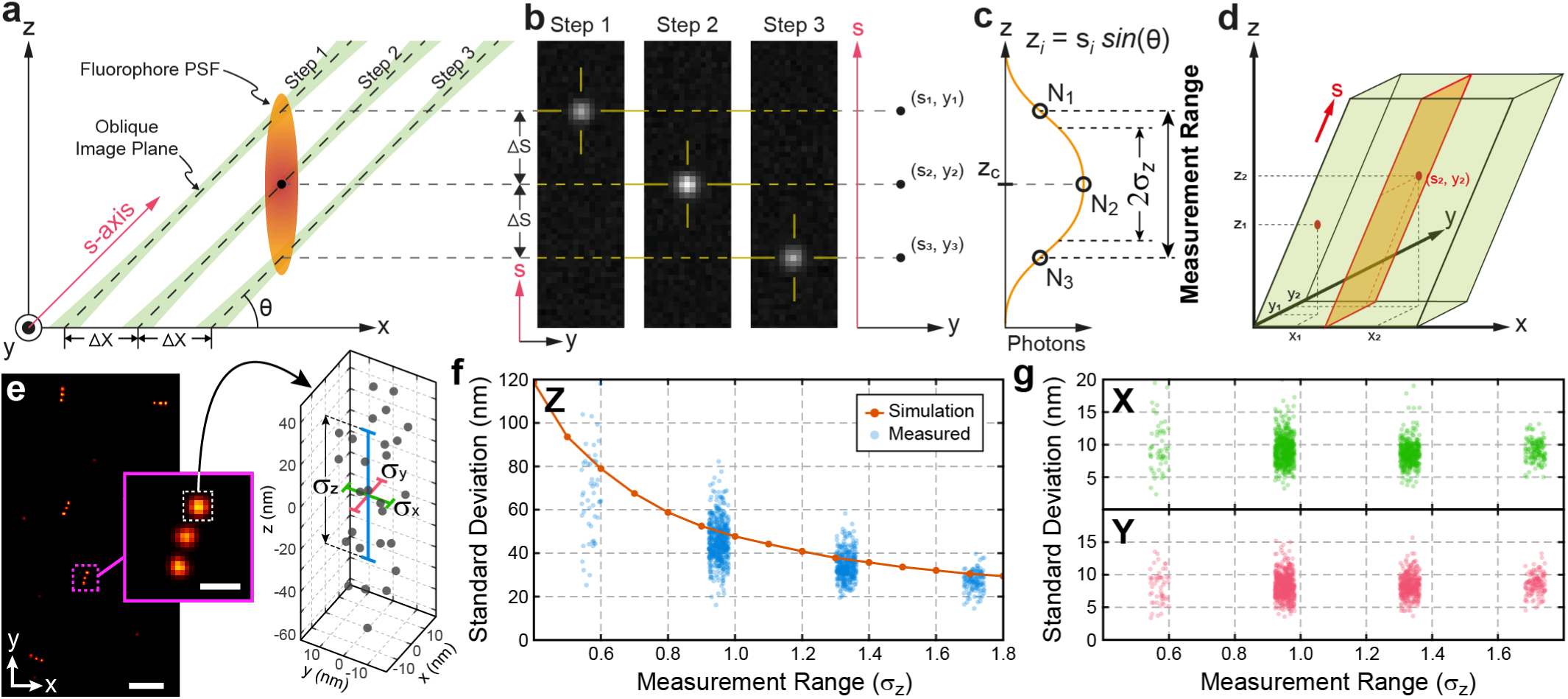
Combining oblique plane microscopy with localization based super-resolution imaging. **(a)** An oblique excitation light sheet is scanned along the x-dimension through the sample volume in discrete steps with spacing ΔX. The scan step size (ΔX) sets the sampling interval for an emitter’s PSF. **(b)** At each step along the x-axis, an image of the PSF is captured on the oblique (s, y) image plane and the emitter is localized to extract the (s, y) coordinates and the number of photons (N) resulting in a set of measurements (s, y, N) for that single emitter. The emitter position is therefore sampled multiple times as the oblique (s, y) image plane is stepped through the sample volume. **(c)** An estimate of the emitter’s z-position is obtained based on the number of photons collected at each s-position. Thus, the s-coordinate, combined with the number of photons at each measurement, encode the emitter z-position. **(d)** The dimensions of the oblique light-sheet determine the extent of the volume along the Z and Y dimensions while the number of steps determine the extent of the volume along the X dimension. During a scan, emitters are localized on the oblique (s, y) plane and their cartesian (x, y, z) coordinates are obtained throughout the entire volume without the need for stitching. Due to the oblique view, the final emitter coordinates are contained in a volume that has a tilted boundary matching the light-sheet angle. (**e**) XY view of DNA-PAINT origami nanorulers with 80 nm linear spacing between marks. Inset shows a magnified view of a single nanoruler with its characteristic three linear marks. Point cloud representations of these individual marks provide a straightforward means of assessing localization precision by calculating the standard deviation (σ) along the X, Y and Z dimensions. (**f**) Measured Z standard deviations of multiple individual nanoruler marks, together with simulation data, as a function of measurement range. The measurement range is normalized to the standard deviation of the gaussian distribution shown in **(c)** and is expressed in units of (σ_z_). For our system σ_z_ = 256 nm, as determined experimentally. (**g**) Measured X and Y standard deviations of multiple individual nanoruler marks as a function of measurement range. Scale bars: 500 nm **(e)**, 100 nm for magnified view in **(e)**.

### SMLM over extended sample volumes

To demonstrate the ability of Intrinsic3D to capture large volumes with high resolution, we imaged the endogenous nuclear pore complex protein NUP96 tagged with GFP in U2OS cells using DNA-PAINT with an anti-GFP-nanobody conjugated to a docking strand (**Fig. 2a**).

**Figure 2.**
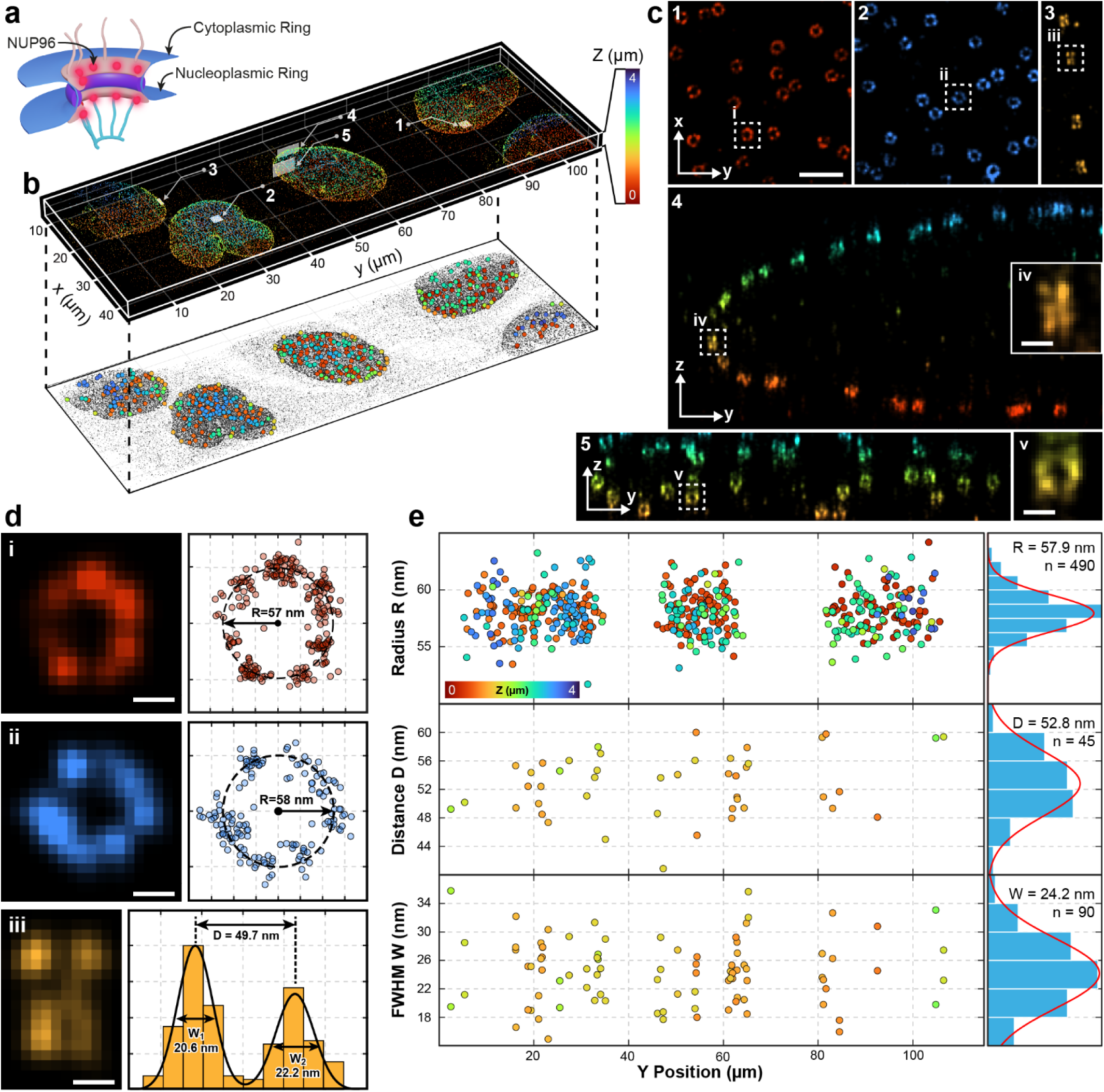
Volumetric imaging of U2OS cell nuclei. **(a)** Cross section schematic of an NPC showing the nucleoplasmic and cytoplasmic rings and the location of the NUP96 protein. (**b**) 3D reconstruction of a ∼ 40 x 115 x 4 µm^3^ (XYZ) volume capturing several U2OS cell nuclei (top) and an XY projection (bottom) showing the locations of selected NPCs used in subsequent analysis in **(e)**. NPC markers are color-coded for depth. Five planes are highlighted in the 3D reconstruction (**b, top**) corresponding to the panels of **(c)**. **(c) panels 1** and **2** show magnified XY views of NPCs from the bottom and top, respectively, of cell nuclei where individual NPC rings are resolved. **Panel 3** shows an XY view at the equator of a nucleus where cross sections of NPCs are obtained and the separation between the cytoplasmic and nuclear rings can be measured. **Panel 4** displays a YZ cross-sectional view of a nucleus where side views of the NPC along the YZ plane are obtained. Here, the separation between the nucleoplasmic and cytoplasmic rings is visible only along the Y dimension (**inset iv**). **Panel 5** shows the YZ face of a nucleus where individual NPCs are resolved as rings also shown in **inset (v)**. Boxed regions in **panels 1-3** are used in **(d)** for example analysis. **(d)** Boxed regions from **(c)** are shown with corresponding example fits. **(e)** Plots showing measured radius (R = 57.9 ± 1.6 nm, n = 490), distance between the nuclear and cytoplasmic rings (D = 52.8 ± 4.54 nm n = 45) and fitted FWHM of each NPC profile (W = 24.2 ± 4.92 nm n = 90) as a function of the Y position. Markers are color-coded for depth corresponding to (b). Scale bars: 500 nm **(c, panels 1-5, shown in panel 1 only)**, 100 nm **(c, iv, v),** 50 nm **(d)**. Images are representative of 5 independent imaging experiments.

We imaged oblique volumes extending ∼**30 x 110 x 15** µm^3^ in X and Y and S dimensions, respectively, capturing whole nuclei of multiple cells (**Fig. 2b, Supplementary Movie 1**). The characteristic NPC ring structure is easily observed on both the top and bottom surfaces of all imaged nuclei (**Fig. 2c, panels 1, 2** and **Supplementary Movie 2**). Similarly, cross sectional views of the nuclei reveal the ∼50 nm separation between the cytoplasmic and nuclear rings. (**Fig. 2c, panel 3**). YZ cross-sections resolved the cytoplasmic and nuclear rings along the Y axis but not along the Z axis **(Fig. 2c, panel 4, inset iv)**, consistent with the expected ∼3.5-fold lower axial resolution **(Supplementary Fig. 2)**. Despite this reduced axial resolution, YZ cross-sectional views also resolved individual NPC rings **(Fig. 2c, panel 5, inset v)**.

Because the cell nuclei span the entire volume, they can be used to quantify performance across different regions of the volume by measuring the NPC radius, the separation between the cytoplasmic and nucleoplasmic rings of NPCs in cross section, and the FWHM of the ring cross sections as a proxy for resolution (**Fig. 2d, i, ii, iii**). NPCs were selected across the whole volume (**Fig. 2b**, lower projection) and fit to extract the radius (**Fig. 2e, upper plot**), the cross sectional cytoplasmic and nuclear ring separation (**Fig. 2e, middle plot**) and the widths (FWHMs) of the two cross-sectional ring features (**Fig. 2e, lower plot**). No change is seen in the radius, cross section separation, or FWHM over the imaged volume indicating consistent resolution throughout the volume. Pooling all selected NPCs, the average radius was measured as **57.9 ± 1.6** nm (mean ± std, n = 490) and **52.8 ± 4.54** nm (mean ± std, n = 45) for the distance between the cytoplasmic and nuclear NPC rings (**Fig. 2e**) in good agreement with previous work ^24^. Furthermore, the FWHM of each gaussian provides an experimental measurement of the resolution along the XY dimensions with an average value of **24.2 ± 4.92** nm (mean ± std, n = 90) (**Fig. 2e**).

### Expediting 3D imaging

Previous volumetric image techniques are slow because thousands of frames covering partially overlapping regions at each z-step are required to cover a complete volume, followed by additional post-processing to stitch data from multiple sub-volumes into a complete 3D dataset. Intrinsic3D images large volumes faster because each scan captures a complete volume and 3D emitter positions are measured directly in the complete imaging space with minimal post-processing or the need for stitching. Like previous OPM implementations, the extent of the light sheet in the S and Y dimensions determines the final volume’s Z depth and Y dimension, respectively, while the X dimension is set by the scan range (**Fig. 1**). Given that the PSF size along the X dimension is ∼300 nm, a step size of ∼150 nm would provide Nyquist sampling along the X dimension. However, in our implementation the emitter is localized on the Y-S plane, and the lateral localization of an emitter is independent of the X step size. Rather, the X step size effectively sets the PSF’s Z sampling interval (**Methods**). Because a minimum of two PSF sampling points is sufficient to localize an emitter’s position in Z, larger steps in the 500-700 nm range can be used. Using a scan step size well beyond Nyquist sampling dramatically accelerates data acquisition.

To demonstrate fast volume acquisition, we imaged NPCs in stage **6-8** *Drosophila* egg chambers. Specifically, we imaged endogenously-tagged NUP160-Halo labelled with an anti-Halo nanobody conjugated to a docking strand^16,25^. *Drosophila* egg chambers feature large interior nurse cells surrounded by a curved monolayer of smaller follicle cells (**Fig. 3a**). This sample geometry allowed us to image multiple complete follicle cell nuclei at different depths and portions of the larger nurse cell nuclei (**Fig. 3b-e, Supplementary Movie 3, Supplementary Movie 4**). Because NPCs have a well-defined geometry, we reasoned that tracking their appearance, as localizations accumulated during imaging, would allow us to estimate the acquisition time required to reconstruct NPCs throughout the volume (**Fig. 3f**). We captured ∼**20,000** oblique volumes of ∼**20** x **90** x **40** µm^3^ (XYS) with a **700 nm** scan step size and a **20 ms** exposure time per step. These settings allowed us to reach sample depths of ∼**14** µm with an imaging speed of **1.67** volumes per second and a total acquisition time of ∼**200** min.

**Figure 3:**
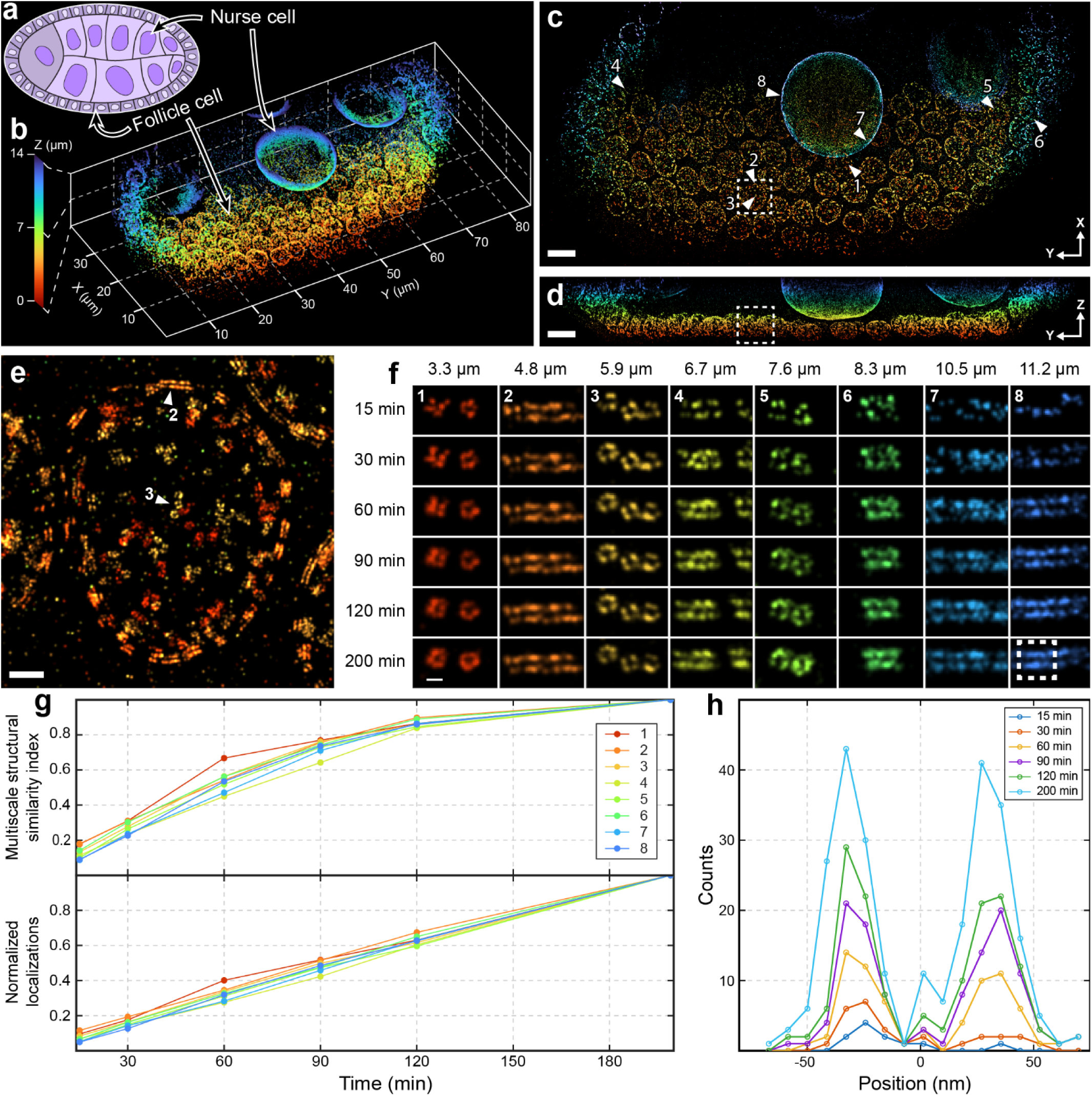
Quantifying NPC appearance during acquisition to evaluate imaging speed. **(a)** Schematic of a *Drosophila* egg chamber showing the follicular epithelium and nurse cells. (**b**) 3D reconstruction of a ∼**45 x 90 x 15** µm^3^ (XYZ) volume capturing multiple follicle cell nuclei along with portions of the much larger nurse cell nuclei located deeper in a stage 7 *Drosophila* egg chamber. (**c**) XY projection of the 3D volume, colour-coded by depth, showing the locations of the NPCs used to assess imaging performance at different time points. Scale bar: 10 µm (**d**) YZ projection, color-coded for depth, of the 3D volume. Scale bar: 10 µm. The boxed region in (**c**) and (**d**) is used for (**e**), an XY projection of a single follicle cell nucleus color-coded for depth. NPCs on the basal side appear as red circles, while NPCs on the apical side appear as yellow circles. The paired traces outlining the nucleus correspond to nucleoplasmic and cytoplasmic NPC rings as visualized in side-view projections. Scale bar: 500 nm (**f**) Montage images showing the evolution of the NPC signal from the numbered regions of the volume shown in (**c**) over the course of imaging. Scale bar 100 nm. (**g**) Multiscale structural similarity index comparing the NPC images shown in (**f**) with their corresponding final NPC images. The number of localizations in each image shown in (**f**) normalized to the corresponding total number of localizations accumulated over the full imaging duration are also shown at the different timepoints (**h**) Evolution of the cytoplasmic and nucleoplasmic NPC ring signals for the portion of the nurse cell shown in (**f**). Images are representative of 5 independent imaging experiments.

To evaluate NPC progression with time, NPCs at a variety of depths were selected (**Fig. 3c-e**) and their images were reconstructed at 30-minute intervals (**Fig. 3f**), where each image included all localizations up to that time point. Selected NPCs included both front views from the tops and bottoms of cell nuclei where NPCs appear as a ring (**Fig. 3f** **panels 1, 3, 5**) and cross-sectional views from the cell nuclei equator where NPCs appear as paired traces (**Fig. 3f** **panels 2, 4, 6, 7, 8**). At each timepoint, we measured the structural similarity index (SSIM) of these reconstructions relative to the final image, which was generated using all localizations accumulated during the full course of imaging (**Fig. 3g**). NPC traces become detectable within the first 30 minutes, and most structural features were resolved after 90 minutes (**Fig. 3f**). Beyond 90 minutes, the number of localizations continues to increase throughout the volume, enhancing the contrast and brightness of the reconstructed image (**Fig. 3g, h**). A more complete representation of the final image (**200** min) is achieved after ∼**120** minutes, reaching a SSIM above 0.8 (**Fig. 3g**). Expanding the volume along the Y and Z dimensions comes at no additional cost in acquisition time, and it is only the length of the scan and the step size along the X dimension that impact data collection times. A complete timelapse of the ROIs found in **Fig. 3f** is available in **Supplementary Movie 5**.

### Investigating protein organization and domain allocation in whole cells

To highlight the utility of imaging multiple whole cells simultaneously with Intrinsic3D, we examined the localisation of the homophilic adhesion molecule, E-cadherin in the *Drosophila* follicular epithelium^26^. Although E-cadherin organization has been investigated using several super resolution techniques, only one study using 2D DNA-PAINT resolved the separation between opposing E-cadherin complexes on adjacent cell membranes ^27^. We imaged endogenous E-cadherin tagged on its C-terminus with GFP^28^ in a volume of 30 x 100 x 7 µm^3^ (x, y, z) in a stage 7 *Drosophila* egg chamber, capturing many entire follicle cells **(Fig. 4a(i))**. To analyse the E-cadherin distribution, we segmented three complete cells **(Fig. 4a(ii) and Supplementary Movie 6)** into adherens junction, apical membrane, cytoplasm and lateral membrane, **(4b(i-v) and Supplementary Movies 7 and 8).** The relative amounts of E-cadherin in the four segmented domains were measured for each cell, correcting for the fact that half the E-cadherin in the spot and adherens junctions comes from the neighbouring cells. This revealed that there is significantly more E-cadherin in lateral spot adherens junctions (spot AJs) than the adherens junction **(Fig. 4c).** To analyse the distribution of spot adherens junctions, we built a model of the lateral membrane (**Fig. 4c** **and Supplementary Movie 9**) and unwrapped the lateral membrane on to a 2D plane and marked the positions of the vertices where three cells meet (**Fig. 4b(vi), 4d**). This revealed that the spot adherens junctions **(Fig. 4e)** are enriched near these tricellular junctions **(Fig 4f)**, which are presumably sites of higher mechanical stress. This enrichment and the observation that there is more E-cadherin in spot adherens junctions than adherens junctions suggest that the former play a more important role in cell-cell adhesion than previously appreciated.

**Figure 4.**
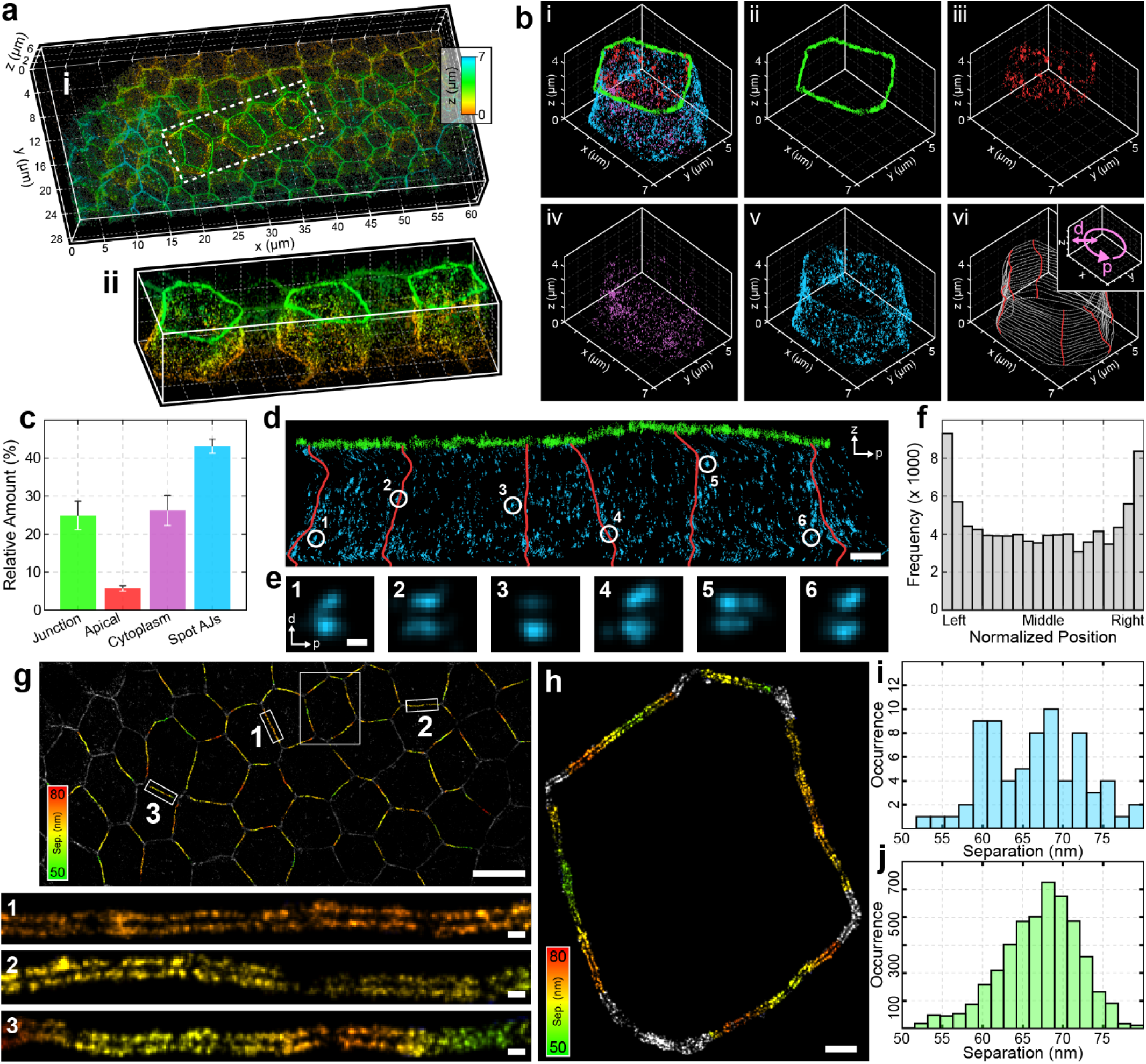
Intrinsic3D whole cell imaging of E-cadherin. **(a)** GFP tagged E-cadherin imaged with Cy3B DNA-PAINT in the follicular epithelial cells of a stage 7 *Drosophila* egg chamber **(i)**. The boxed region in **(i)** highlights the three cells segmented in **(ii)** and used for analysis. The rightmost cell in **(ii)** is used for **(b, d, e, and h). (b, i)** All three cells were further segmented into four domains with the right most cell shown as an example: **(ii)** apical adherens junction (green), **(iii)** apical membrane (red), **(iv)** cytoplasm (purple) and **(v)** spot AJs (lateral membrane, blue). The relative amounts of E-cadherin in the four domains (lateral, junction, apical, cytoplasmic) for all three cells were tabulated and shown in **(c)** with error bars as ± the standard deviation. A model of the lateral membrane was generated as shown in **(b, vi)** and used to transform (x, y) lateral membrane localizations to Frenet coordinates (p, d) as diagrammed in the insert to **(b, vi)**. The complete lateral membrane and junction were unwrapped onto a 2D (p, z) plane as shown in **(d)** with tricellular junctions (vertices) indicated as red lines. Spot AJs were selected with examples indicated in **(d)** and corresponding (p, d) images shown in **(e)**. The position of each localization on the lateral membrane, relative to its two bounding left-right vertices, was found and a histogram of normalized positions is shown in **(f)**. The peaks at the left and right sides suggest that E-cadherin is enriched close to tricellular junctions. Adherens junctions from **(a)** were segmented and fit with a Gaussian mixture model to determine the separation between opposing cell membranes along each junction. Fitted separation values were used to generate a junction map color-coded by membrane separation **(g)** with magnified examples boxed in white. The junction of the cell used in **(b-d)** is shown in **(h)** colour-coded for E-cadherin separation. Selected Spot AJs from the three segmented cells were fit with a two peak 2D Gaussians to determine the separation with the resulting distribution shown in **(i)** and mean value of 67.3 ± 7.3 nm (mean ± st. dev., N = 75). The distribution of fitted adherens junction separation is shown in **(j)** with a mean of 67.1 ± 4.95 (mean ± st. dev., N = 3616 sampling points). Scale bars: (d) 1 µm, (e) 50 nm, (g) 5 µm, (g, 1, 2, 3) 100 nm, (h) 500 nm.

E-cadherin complexes in the membranes of adjacent cells were well-resolved, with a clear space between them (**Fig. 4g and h)**. The distance between E-cadherin complexes on either side of the junction varied between cells, junctions and within the same junction as shown by the colour coding in (**Fig. 4g, 1-3 and h).** The mean distance was 67.1 ± 4.95 (mean ± std, n = 3616 sampling points), which is larger than the distance measured using 2D PAINT in the *Drosophila* embryo (∼ 42 -55 nm), which imaged the same C-terminal GFP tag^23^ . This may be in part due to the fact that the 2D data include junctions that are at an angle to the imaging plane, leading to an under-estimate of this distance, whereas all junctions were measured in the correct plane in our 3D data. However, the follicular epithelium is subjected to expansive forces from the growing germline cyst that it surrounds, and the coupling of E-cadherin to the actin cytoskeleton to resist this force may stretch E-cadherin, so that the C-termini of molecules engaged in homophilic adhesion are pulled further apart. The distance between the E-cadherin complexes in opposing membranes for spot adherens junctions was 67.3 ± 7.3 nm (mean ± std, N = 75), which is not significantly different to that of the adherens junctions, suggesting that they are subjected to similar mechanical forces. Distributions for spot junctions and junctions are shown in **Fig. 4i** and **Fig. 4j**, respectively. Our investigation of E-cadherin distribution illustrates how volumetric imaging of whole cells with Intrinsic3D enables novel visualization and analysis approaches.

### Investigating junction architecture using 2-colour 3D SMLM

To demonstrate the compatibility of Intrinsic3D with multicolour imaging modalities, we performed exchange PAINT to investigate the organization of E-cadherin in the adherens junctions and Gliotactin^29^, a transmembrane protein that is a component of the tricellular septate junctions^30^ that restricts paracellular diffusion where three cells meet. As septate junctions only mature later in egg chamber development, and Gliotactin is only expressed in follicle cells from stage 10b of oogenesis onwards^31,32^, we imaged follicle cells in stage 12 egg chambers (**Fig. 5a**, **Supplementary Movie 10).** Higher magnification views show that Gliotactin forms string-like structures that are mainly parallel to the plane of the epithelium around the tricellular junction **Fig. 5b-f).** There is a gap between the adherens junction belt and the spot adherens junctions, which is filled with Gliotactin at the tricellular junctions (asterisks in **Fig. 5** **c and e**). This gap is not observed in younger egg chambers in which the septate junctions have not yet formed, strongly suggesting that the septate junctions exclude spot adherens junctions from this region.

**Figure 5:**
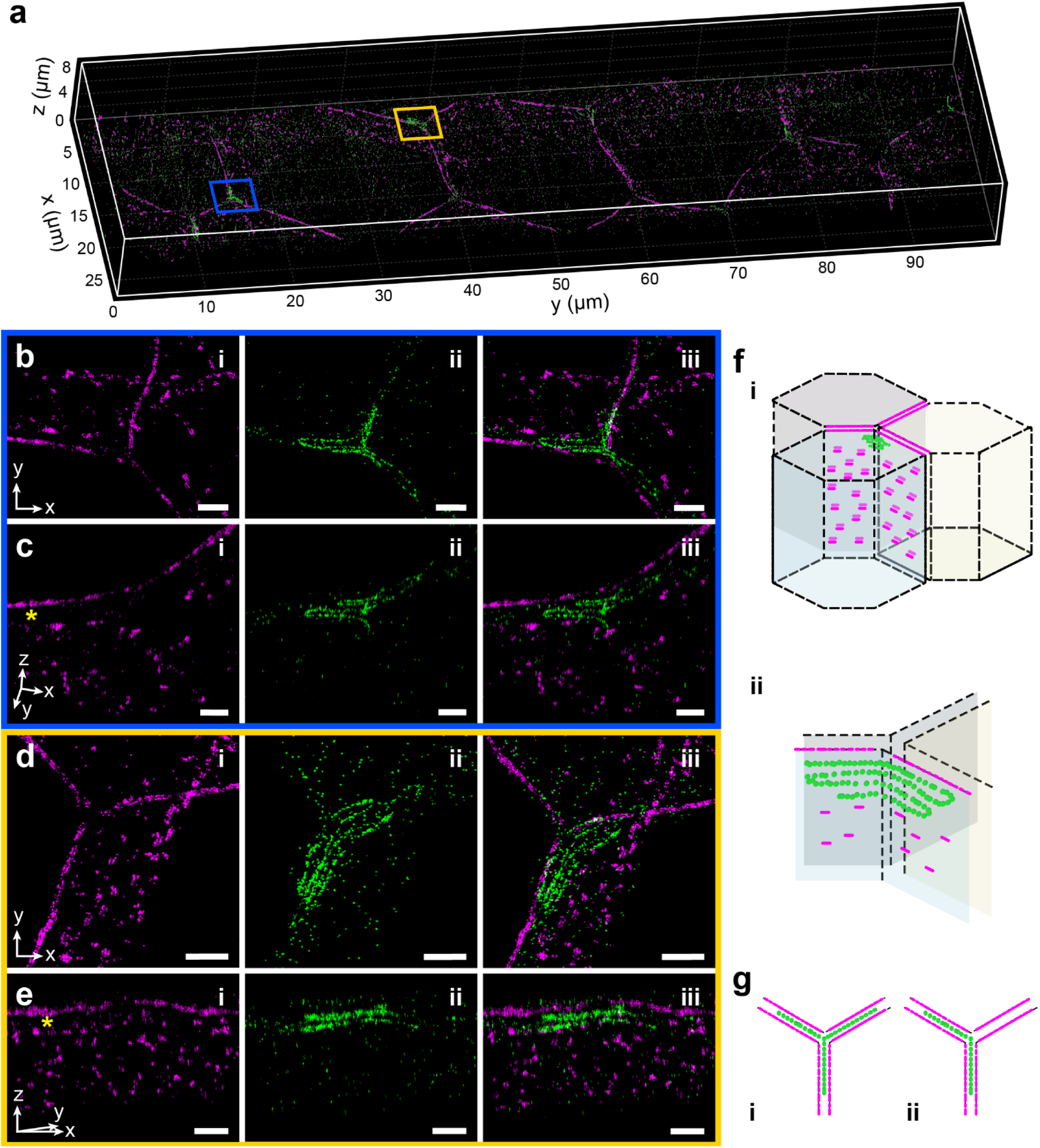
E-Cadherin and Gliotactin organisation in follicle cells. **(a)** 3D reconstruction of a 30 x 100 x 9 µm (XYZ) volume of a stage 12 *Drosophila* egg chamber, showing E-Cadherin-3x-mKate (magenta) and Gliotactin-YFP (green) in the follicular epithelium. (**b**) XY projection of the blue boxed region from **(a)** showing the tricellular junction with **(b, i)** E-Cadherin, **(b, ii)** Gliotactin **(b, iii)** merge. **(c)** View normal to the lateral membrane of two of the cells from the blue boxed region from **(a)** showing the protein organization on the lateral membrane, **(c, i)** E-Cadherin, **(c, ii)** Gliotactin **(c, iii)** Merge. Asterisk in **(c, i)** indicates gap between the adherens junction and the spot adherens junctions. (**d**) XY projection of the yellow boxed region from **(a)** showing the tricellular junction **(d, i)** E-Cadherin, **(d, ii)** Gliotactin **(d, iii)** merge. **(e)** Normal view of the lateral membrane of two of the cells from the yellow boxed region in **(a)** showing the protein organization on the lateral membrane, **(e, i)** E-Cadherin, **(e, ii)** Gliotactin **(e, iii)** merge. Asterisk in **(e, i)** indicates gap between the adherens junction and the spot adherens junctions. **(f)** Diagram of E-Cadherin and Gliotactin at the tricellular junctions **(i)** with a magnified view shown in **(ii)** with the proteins shown in only one of the three cells. **(g)** Diagram of Gliotactin in tricellular junctions where it is present at all three interfaces **(g, i)** and present at only two interfaces **(g, ii)**. All scale bars are 1 µm.

In some cases, Gliotactin is present on all three bicellular interfaces around the tricellular junction, forming a complex folded string structure (**Fig. 5b**, **c** and **Supplementary Movie 11**). In other junctions, the Gliotactin “string” is only present on two of the three interfaces (**Fig. 5e**, **f, g** and **Supplementary Movie 12**). The reason for this variability in tricellular junction structure is not clear, but 20/34 tricellular junctions had Gliotactin at only two interfaces, and 14 at all three.

## Discussion

In this work we introduce Intrinsic3D, a method that combines oblique light-sheet microscopy with SMLM to localize emitters in 3D within large volumes. Intrinsic3D relies on a fast-scanning oblique light sheet to sample an emitter’s PSF at multiple locations during a single blinking event. Because the position of the light sheet within the full volume is known at each PSF sampling step, the photon plus light sheet position information is combined to unambiguously determine an emitter’s Z position within the volume. This does not require any z position calibration, or PSF engineering, and avoids the need to stitch sub volumes together in post processing. Because redundant imaging of over-lapping neighbouring sub-volumes is not necessary, our method reduces imaging time and simplifies post-processing. Intrinsic3D localizes emitters on the oblique plane. This localization geometry removes the constraint of Nyquist sampling along the scan dimension and allows for larger scan steps. In fact, simulations and experimental data with DNA origami show that larger scan steps (PSF measurement range) improve z localization. Without the requirement to Nyquist sample in the scanning dimension, the limit on how big the scan step size can be is imposed by the detectable PSF size itself. If the step size is too big, the PSF will only be sampled once, and the emitter’s z-position cannot be localized. The maximum step size is set by the axial size of the PSF, the NA of the light-sheet and the angle of the oblique imaging plane. In the work presented here, the oblique plane is set at 26°, the PSF’s axial FWHM is ∼800 nm and the NA of the excitation light-sheet is ∼0.14. Under these conditions, a scan step of ∼700 nm along the x-axis is sufficient to ensure a minimum of 2 measurements on the emitter position for z localization. This represents a seven-fold increase in speed, compared to Nyquist sampling with steps of ∼100 nm along the X dimension. The depth (z) and width (y) dimensions of the final volume are determined by the extent of the region illuminated by the light sheet and how much of that region is captured by the camera. Thus, changing the width and depth of the imaged volume by changing the light sheet size and/or camera ROI has no impact on data acquisition time.

The confocal parameter (length) of the light-sheet sets the maximum achievable depth while the light sheet thickness controls the optical sectioning. Here, a gaussian light sheet with an NA of 0.14, a length of ∼40 µm and thickness of ∼1 µm^33^ was used. Imaging volumes with larger depths could be achieved by employing lower NA light-sheets that have longer lengths but at the cost of reduced optical sectioning.

We note that instead of imaging an oblique plane, a plane perpendicular to the optical axis of the primary objective could be imaged. This would set the oblique angle to zero and the optical setup would assume the configuration of a remote focusing system (**Supplementary Fig. 3**) ^34^. This configuration maximizes the volume along the X and Y dimensions but scanning along the Z dimension would have to be performed by stepping the tertiary objective (**Supplementary Fig. 3**). Additionally, this configuration would require widefield illumination that does not provide any optical sectioning. Nonetheless this could be a viable option for certain imaging applications with limited depth, such as cultured cells, if the lack of optical sectioning could be compensated for by using fluorogenic and/or optimized imager probes that reduce the background in PAINT imaging ^35,36^.

We have demonstrated that our method is well-suited for PAINT imaging where the blinks are produced by transient binding events with durations ranging between 0.2-1 sec. However, since the camera in our setup supports exposure times down to 2.5 ms, this method can also be implemented with other variants of SMLM including STORM, where the duration of photophysical blinking can be tuned in the 50-100 ms regime by adjusting the excitation intensity ^37^.

Additionally, combining the optical sectioning of OPM with SMLM has the added benefit of background suppression when imaging deeper into 3D specimens away from the cover glass. Although fluorogenic probes that reduce background from freely diffusing emitters have recently been introduced, taking advantage of the optical sectioning offered by OPM ensures that the full suite of probes, fluorogenic or not, may be used which may be attractive as more multiplex imaging schemes are introduced. As demonstrated here, high-quality 3D imaging over large tissues volumes is possible with non-fluorogenic probes and providing new insights into protein localisation in cells.

Importantly, Intrinsic3D is fully compatible with recent advances in multiplexed and quantitative imaging. Combining these methods with Intrinsic3D would increase throughput and enable spatial proteomics in tissue samples with nanometre level resolution in 3D and single protein sensitivity.

## Methods

### Microscope Setup

The OPM system used for data collection was integrated into a commercially available inverted microscope base (Olympus, IX81) with the help of a scan-mirror pair. The system was identical to the one described previously^38^ with minor modifications to allow for Cy3B imaging (**Supplementary Fig. 3**). These include the incorporation of a laser line at 546 nm, the addition of a dichroic mirror (Chroma, 59007bs) to separate Cy3B fluorescence from the excitation laser and an additional band pass filter (Chroma, 59007m).

### Data Acquisition

Data acquisition and instrument control were performed with an NI field programable gate array (FPGA) data acquisition card (DAQ) (NI, PCIe-7852R) and custom control software developed in the LabView environment paired with an existing control package publicly available at (https://github.com/Gurdon-Super-Res-Lab/Microscope-Control). Unlike traditional imaging modalities, the depth of the acquired volume in OPM is set by the size of the camera (Hamamatsu, Orca-Flash 4.0 v3) ROI. This is because the camera is viewing an oblique plane and the size of the ROI along the s-dimension, sets the imaging depth. The other dimension of the ROI sets the y-size of the final volume, and the scan length sets the x-dimension. Thus, the ROI size is set based on the sample and required imaging depth (**Fig. 1d**). After setting an ROI on the camera, the scan length and step size were set. The FPGA card was used to synchronize mirror stepping and camera acquisition. This card has both digital inputs and outputs and analogue voltage outputs. The camera sent a trigger ready signal to the FPGA card which, in turn, triggered one mirror step. After the mirror step was complete, the FPGA card triggered the camera to capture the next frame. This continued until the complete volume had been scanned at which point the FPGA card returned the mirrors to their starting positions and the process was repeated. Data were acquired in blocks (cycles) of 70 to 100 volumes, taking ∼ 1 minute to acquire. The z-position of the primary objective lens was continuously monitored, and the voltage ratio of the scanning mirrors was adjusted based on a previous calibration to compensate for axial lens motion. Without this correction, axially translating the objective lens will misalign the back aperture and the focused excitation spot causing the light-sheet angle to change while scanning. Each image file containing 100 volumes was automatically transferred to a network attached storage (NAS) drive for processing. Data acquisition times for single colour imaging typically ranged between 2 and 4 hours depending on the scan length, number of steps and camera exposure time. The camera had an effective pixel size of 117 nm along the s- and y-axis.

### Data Analysis

Raw camera frames were converted from ADUs to photons based on manufacturer supplied gain information and experimentally determined offset. Candidate single molecule events (blinks) were identified using a wavelet-based method implemented in MATLAB ^39^. After candidate locations were identified, 7 x 7 subregions were cropped for each candidate location and fit using an MLE fitting package for 2D Gaussians^40^. This provided a set of (y, s) coordinates for each detected blink along with the number of photons.

During acquisition, a single blink event is sampled multiple times at several sequential mirror positions (steps) as the excitation light-sheet and detection plane are scanned through the sample volume. This corresponds to a set of camera frames where a single blink is recorded multiple times with slightly shifted s-positions (s-step) in each sequential camera frame (proportional to the mirror step size). Thus, each (y, s) coordinate must be clustered with the other (y, s) coordinates arising from the same blinking event in subsequent camera frames. When a first (y, s) event is identified, the subsequent (y, s) coordinates will always appear one s-step away and always proceed in the same direction. This makes clustering (y, s) coordinates into PSF tracks straightforward as the stepping direction and s-step size are already known. Each time an unclustered (y, s) coordinate is identified, the subsequent frame is checked for a (y, s) coordinate one s-step away in the correct direction. If another coordinate is identified, it is added to the PSF track (cluster), and the next frame is checked. This process is repeated on each subsequent frame until no new (y, s) coordinate, in the positive s-direction, one s-step away, is found. Each (y, s) coordinate associated with the newly identified PSF track (cluster of (y, s) points arising from a single blink event) is tagged with a unique track number. In this way all the (y, s) coordinates within a single volume are clustered into PSF tracks (or excluded if they are not part of a track). Thus, each single molecule blink event is sampled multiple times and represented by a set of (y, s) coordinates from which the (x, y, z) position is estimated **(Supplementary Fig. 4)**.

### Localization in X and Y

The y-position was simply taken as the localization precision weighted average of all the y_i_-positions in a PSF track as 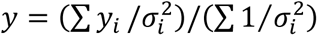. Each time the mirrors were stepped, the x-position of the blink event appears to change due to the excitation light sheet and detection plane positions translating one mirror step along the x-dimension in the sample volume. To correct for this, the x-position for each PSF track was computed as *x_i_* = *s_i_* cos(*θ*) − *n* · ΔX where *n* is the mirror step number (position within the sample volume) and ΔX is the mirror step size. The final x-position was taken as the localization precision weighted average of the *x_i_*′*s* or 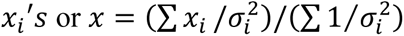.

### Localization in Z

The axial (Z) position of each localization was estimated by performing a weighted least-squares fit to the photon numbers measured at all positions (*s_i_*) along a PSF track. At each position, the number of photons *N_i_* and its associated uncertainty (*σ_Ni_*), as determined from the upstream 2D MLE fit, were used to construct a Gaussian model, 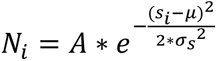 with a fixed width (*σ_s_*), obtained from the experimental PSF of the system, and free parameters, the peak amplitude (*A*) and the expected s position of the molecule (*µ*). To account for differences in measurement uncertainty between positions, both the data and model were scaled by ^1^/*_σNi_* prior to fitting. The fit was performed using the Levenberg–Marquardt algorithm (MATLAB’s lsqcurvefit) and the final Z position of the molecule was calculated as *μ* ∗ sin(*OPM angle*). Localization precision was estimated empirically via parametric bootstrapping: 50 synthetic sets of photon measurements were generated by adding Gaussian noise drawn from (*σ_Ni_*), and each set was refit independently to yield a distribution of *µ* estimates. The standard deviation of this distribution gave the precision along *s*, which was scaled by sin(*OPM angle*) to obtain the final localization precision along Z.

### Simulations

#### Number of samples and sampling range

To characterize how the Z localization precision depends on the number of sampling points (N) and the sampling range, we performed Monte Carlo simulations of Z localization for an emitter with a 1D Gaussian intensity profile centred at the true emitter position, with standard deviation σ (**Supplementary Fig. 1a, b**).

We considered sample counts of N = 2, 4, 8, and 16, and sampling ranges from 0.25σ to 3σ in steps of 0.25σ. For each combination, N positions were uniformly distributed within the detection range, constrained to lie within ±2σ of the emitter position (**Supplementary Fig. 1a, b**). Expected photon counts at each position were computed from the Gaussian profile and corrupted with Poisson noise. For each noisy realization, the emitter’s Z position was estimated via a Gaussian model using maximum likelihood estimation (MLE) under a Poisson counting-noise assumption. Fits that failed to converge were discarded. This procedure was repeated for 10,000 independent noise realizations per (N, range) condition. Localization precision was quantified as the standard deviation of the estimated positions across the 10,000 successful fits (**Supplementary Fig. 1c**). The localization precision improved substantially with increasing measurement range and showed a ∼5x improvement as the measurement range increased from 0.25σ to 1σ. Furthermore, for a fixed measurement range increasing the number of measurements offers an additional but more modest improvement in the localization precision. These results suggest that maximising sampling range is the primary lever for improving axial localisation precision, with the number of measurements playing a secondary role, informing the choice of step size and scan range used in subsequent experiments.

#### Step size, exposure time, scan length

To determine how the step size and fluorophore blinking kinetics jointly affect the recovery of usable localisation tracks, we developed a Monte Carlo simulation of an Intrinsic 3D experiment. The axial point-spread function (PSF) of an emitter was modelled as a Gaussian. Emitters were randomly distributed along the scan axis (density 100/µm), and each was assigned a stochastic emission trace of 1000 sequential binding events, with exponentially distributed "on" and "off" interval durations. For every frame in which a molecule was "on" and within the detection window, an expected photon count was computed from the Gaussian PSF and a Poisson-distributed count drawn from this expectation, explicitly incorporating shot noise into the threshold-based detection decision. Contiguous runs of positive detections were identified as discrete "tracks" and linked back to their underlying binding event, allowing tracks of two or more measurements (the minimum for axial position estimation) to be distinguished from single-frame detections that cannot be localized (**Supplementary Fig. 1d**). Combining these results with the number of missed binding events due to spatiotemporal separation with the light-sheet allowed us to estimate the fraction of localized binding events vs total number of binding events as a function of the time required to complete one full imaging cycle (**Supplementary Fig. 1e**). To facilitate comparison, we normalized the cycle time (t_c_) to the on-time of the fluorophore (t_on_) (0.25-12.25) (**Supplementary Fig. 1e**) and the step size ΔX to the size of the detection-range (D), defined as the range around an emitter’s position within which a binding event produces a detectable number of photons, (0.2-0.8) (**Supplementary Fig. 1f-i**). The fraction of localized binding events fell sharply with increasing cycle time indicating that imaging can be expedited when the time required to complete a full scan of the imaged volume is short relative to the duration of a binding event. Furthermore, step sizes up to ∼ half the detection range (0.5*D) do not substantially increase the proportion of tracks with single measurements that cannot be localized. Taken together these data suggest that larger step sizes up to ∼half the detection window expedite imaging as they reduce scan time. This comes with no significant cost in resolution as larger steps also extend the sampled range of the emitter PSF improving axial localization precision.

### Drift Correction

Nearest-paired cloud (NP-Cloud) drift correction developed by Hou *et al.* ^41^ was applied to all images in post-processing. Specifically, after all DCIMG files were fit and all (y, s) coordinates converted to (x, y, z) coordinates, the number of neighbouring points within an approximately 2*σ_xy_* radius by 2*σ_z_* tall volume were found for each localization. Any localization that had 0 or 1 neighbours within the search volume was discarded. The remaining (x, y, z) coordinates were drift corrected with NP-Cloud and saved without further processing.

### Lateral membrane unwrapping

Custom analysis scripts were developed in the MatLab environment to build a skeleton of the lateral membrane and junction and convert (x, y, z) coordinates to Frenet (p, d) coordinates. Building the lateral membrane skeleton was done with sequentially with overlapping z-slices. Specifically, a single 300 nm thick z-slice of lateral membrane localizations was segmented and converted to polar coordinates (φ, rho). A random subset of points spanning 0 to 2π was used to interpolate rho values at 2000 evenly spaced φ values from 0 to 2 π. This was repeated 20 times for 20 different sets of random points. Each set of interpolated rho values represents a random path tracing the junction. The 20 random paths were averaged to make a single average path that traced the lateral membrane for that z position. The z position was then incremented by 100 nm, a new 300 nm z-slice was segmented and the process repeated until the average path at each 100 nm step was computed. In this way, a frame or skeleton of the lateral membrane was created as shown in Fig. 4b(vi) and used to convert the (x, y, z) localizations of the lateral membrane and junction to Frenet Coordinates. Frenet coordinates use a fixed path as a reference and measure the perpendicular distance between a point and the path (coordinate “d”) and the distance along the path (coordinate “p”) as illustrated in the insert to Fig. 4b(vi). Each path in the skeleton was converted from a set of positions to a set of distances from the origin (φ = 0). For each (x, y, z) localization, the skeleton was used to interpolate the distance along the path from φ = 0 and the distance of the point from the path for that point’s specific z position. In this way, each localization’s (x, y, z) coordinates on the lateral membrane was converted to (p, d, z) and the lateral membrane was unwrapped onto a 2D plane (the p, z plane, similar to an XZ or YZ view) as shown in **Fig. 4c** (blue) along with the junction (**Fig. 4c**, green).

### E-Cadherin Adherens Junction Segmentation and Fitting

Localizations, as show in **Fig. 4**, were manually cropped in x, y and z to an ROI primarily containing adherens junctions. The (x, y) coordinates were used to reconstruct a 2D (x, y)-only image, which was converted into a binary image using a simple threshold, leaving adherens junctions as the predominant remaining features. Next, each pixel in the binary image was assigned an intensity based on the distance to its nearest nonzero pixel. This, combined with the original binary image, was used with a watershed algorithm and the resulting ridge lines were converted to (x, y) coordinates representing an initial guess at adherens junction locations in (x, y). The initial junction coordinates were used to find all localization within a 230 nm (x, y) distance (ignoring z) for that specific junction. An initial linear path in z was generated from iteratively fitting and localizations more than 400 nm away, in z, were excluded. The remaining localizations were fit with a polynomial in xy and, separately, z to generate an initial 3D path following each junction (Supplementary Fig. 5a). Points more than 300 nm away from the initial path in z, and 117 nm in xy, were excluded. Threshold and filter values were determined empirically based on the experimental data.

Next, each set of segmented junction localizations was fit with a spline in xy following its centre. The central spline was used to convert junction (x, y) localization coordinates to Frenet coordinates (arc length and distance from centre, s and d, respectively) creating a “straightened” junction where the centre of the junction was always at d = 0. After straightening, local rotation ("roll") of the junction was corrected. That is, correction was applied in areas where the membrane orientation was not parallel with the z axis. More specifically, local out-of-plane membrane roll was corrected by constructing a smooth centre line and associated local orthonormal frame from the localization data. In sliding windows along the junction, the orientation within the plane normal to the centre line that best resolved the two adjacent populations was estimated using a kurtosis-based projection-pursuit metric. Roll-angle estimates were confidence-weighted, spatially smoothed using circular statistics, and used to rotate localizations into a common reference frame, thereby maximizing separation of the opposing membrane signals along the entire junction. Sparse outliers were excluded before model fitting.

After straightening and correcting for roll, adherens junctions were model as a parallel double-line structure using a Gaussian mixture model and fit using an expectation-maximization (EM). The model represented the adherens junctions as a centre line c(x) and a position-dependent separation s(x). Both c(x) and s(x) were represented using cubic regression splines in truncated power basis form, consisting of a global cubic polynomial and additional truncated cubic basis functions centred at interior knots. Localizations were iteratively assigned to one of the two lines or an optional background component based on Gaussian likelihoods. During the E-step, posterior assignment probabilities (responsibilities) were calculated for each localization from the relative likelihoods of the two curve components and the background component. During the M-step, the coefficients of c(x) and s(x) were jointly updated by weighted least-squares optimization, with the responsibilities providing the statistical weights for each localization. Ridge regularization was applied to suppress overfitting, and linear inequality constraints were used to enforce physically meaningful bounds of 40 to 90 nm. The final model provided estimates of the centre line shape c(x), the position-dependent separation s(x), mixture fractions, background fraction, and point-wise assignment probabilities for each localization (Supplementary Fig. 5b). Convergence was determined from the change in log-likelihood between iterations.

### GATTA-PAINT nanoruler imaging

Oblique volumes were acquired using the GATTA-PAINT nanoruler (GATTAquant HiRes 80R), as described in the Data Acquisition section, with step sizes of 300, 500, 700 and 900 nm along the X dimension. For each step size, the same volume was imaged, and the number of acquired volumes was adjusted to maintain an approximately constant total acquisition time.

#### Drosophila melanogaster lines

Standard procedures were used for *Drosophila* maintenance and experimentation.

E-Cadherin images were taken using the GFP knock-in lines of Huang *et al.*^28^. Freshly eclosed homozygous flies were fattened on yeast for 1-2 days at 25°C before dissection.

Nup160 images were taken using the Nup160-Halo CRISPR line from Cheng *et al.*^16^. Freshly eclosed flies were fattened on yeast for 1-2 days at 25°C before dissection.

To create two colour images of E-Cadherin and Gliotactin we crossed E-Cadherin 3x-mKate^42^ with Gliotactin YFP protein trap line^43,44^. Female flies heterozygous for both tagged proteins were selected and fattened on yeast for 2 days at 25°C before dissection.

#### *Drosophila* sample preparation

E-Cadherin-GFP samples

Fattened ovaries were dissected in Schneider’s medium supplemented with insulin (7.5 µg/ml, Merck I9278). Ovarioles were dissected from the surrounding muscle sheath and were fixed with prewarmed (38°C) fixative (4% methanol-free formaldehyde in 0.5x PBS) for 20 minutes at room temperature, with rotation. Samples were washed with PBS for 5 minutes. They were then permeabilised in 0.2% Triton X-100 in PBS for 5 minutes, followed by 2x 5 minute washes in PBS. Samples were incubated in antibody incubation buffer (Massive Photonics) for 30min. Labelling was performed overnight at 4°C with rotation with 1:300 dilution of the GFP-nanobody conjugated to docking strand (Massive Photonics, either Massive Tag-Q Anti-GFP Kit, or Massive Tag-Q FAST Anti-GFP Kit) in antibody incubation buffer. Samples were washed 3x 10 minutes in 1x washing buffer (Massive Photonics), then in PBS + 0.1% Triton. Samples were washed 1x 5 min in PBS and mounted on 8-well chambered coverslips (Ibidi, 80807) that had been treated with Cell-Tak (Corning, 354240). Samples were imaged in 300 µl of imaging solution (Massive Photonics) with 0.5 nM Imager 3 Cy3B, or 0.5 nM F3-LB Cy3B (Massive Photonics) for GFP.

### Nup160-Halo samples

Samples were treated as above but with the Massive Photonics Massive Tag-Q-FAST Anti-Halo Kit, but incubating with a 1:300 dilution of the nanobody for 1 hour at room temperature, followed by an overnight wash in PBS + 0.1% Triton at 4°C.

### E-Cadherin-3x-mKate, Gliotactin-YFP samples

Samples were treated as for E-Cadherin-GFP except younger egg chambers were discarded, and the older stages fixed. Labelling was performed overnight at 4°C with rotation with 1:300 dilution of each of the GFP-nanobody, and TagFP-nanobody conjugated to docking strands (Massive Photonics, Massive Tag-Q FAST Anti-GFP Kit and Massive Tag-Q Anti-TagFP Kit) in antibody incubation buffer. Samples were washed 1x 5 min in PBS, then incubated (with rotation) in a 1:1 dilution of 80 nm gold nanoparticles (Merck 742023) as fiducial markers for two color registration, then washed 1x 5 minutes in PBS. Samples were mounted on pretreated 8-well chambered coverslips (Ibidi 80807). The 8-well chambered coverslips were treated with Cell-Tak (Corning 354240), followed by incubation with a 1:4 dilution of 80 nm gold nanoparticles (Merck 742023) in dH2O for 10 min, then washed with dH2O before sample mounting. Samples were imaged sequentially. E-Cadherin-3x-mKate was imaged with the TagFP nanobody, using 300 µl of imaging solution (Massive Photonics) with 1 nM Imager 3 Cy3B. Gliotactin-YFP was imaged with the GFP nanobody, using 300µl of imaging solution (Massive Photonics) and 1 nM F3-LB Cy3B (Massive Photonics).

### Cell culture, fixation, and staining

U2OS-Nup96-mEGFP cells were a gift from Jonas Ries (EMBL). This cell line was cultured in culture media composed of DMEM GlutaMAX supplemented with 10% foetal bovine serum (FBS) and 1% penicillin-streptomycin (all from Gibco, Thermo Fisher Scientific) at 37°C and 5 % CO2 and was regularly tested for mycoplasma contamination. For imaging, cells were plated onto 8-well glass bottom chambered coverslips (Ibidi, cat. no. 80807) and left to adhere overnight in culture conditions before fixation.

To visualise nuclear pore complexes via Nup96-EGFP, U2OS cells were prepared according to the protocol described in Pleiner *et al*^45^. Briefly, cells were prefixed in transport buffer (TRB: 20 mM HEPES at pH 7.5 with 110 mM KAc, 1 mM EGTA, 250 mM sucrose) with 2.4% formaldehyde (Fisher Scientific) for 30 secs, following which plasma membrane permeabilization was achieved using TRB with 25 µg/mL digitonin for 8 mins on ice. Cells underwent blocking in TRB with 1% w/v bovine serum albumin (BSA) for 2x 5 mins and staining with 100 nM anti-GFP sdAB (MASSIVE-TAG-Q-FAST anti-GFP, Massive Photonics) in the same blocking solution for 30 mins on ice. Cells were further fixed in blocking solution supplemented with 3% (w/v) formaldehyde for 10 mins, washed in blocking solution 2x 5 mins, before nuclear membrane-specific permeabilization in 0.4% Triton X-100 in Phosphate-Buffered Saline (PBS) for 3 mins. Samples were stained again with 50 nM of the same anti-GFP sdAB in blocking solution for 30 mins on ice. Imaging was performed in Massive Photonics DNA-PAINT imaging buffer with 0.5 nM F3LB-Cy3B (Massive Photonics).

## Acknowledgements

This work was supported by a Wellcome Principal Fellowship (224402/Z/21/Z), a BBSRC project grant (BB/V008595/1) and a centre grant support from the Wellcome Trust (24843) to D.St J., the ERC under the European Union’s Horizon Europe research and innovation programme grant 101224877 (EpiRAP) to E.K.P and D.St J.. J.L.-L. is an Oliver Gatty PhD student.

## Author Contributions

D.St J., G.S. and E.S.A. conceived the project. G.S. and E.S.A. developed the method. G.S. and E.S.A. analysed all the data. E.S.A. performed the imaging experiments with the GATTA-PAINT nanoruler. J.H.R. made the fly lines, optimised protocols, prepared and imaged the *Drosophila* samples. J.L.-L. and E.K.P. provided U2OS cell samples and E.S.A. performed the imaging experiments. D.St J. acquired funding and supervised the project. G.S., J.H.R, E.S.A and D.St J. wrote the manuscript with input from all authors. All authors participated in the discussions and data interpretation.

## Supplementary Figures

**Supplementary Figure 1.**
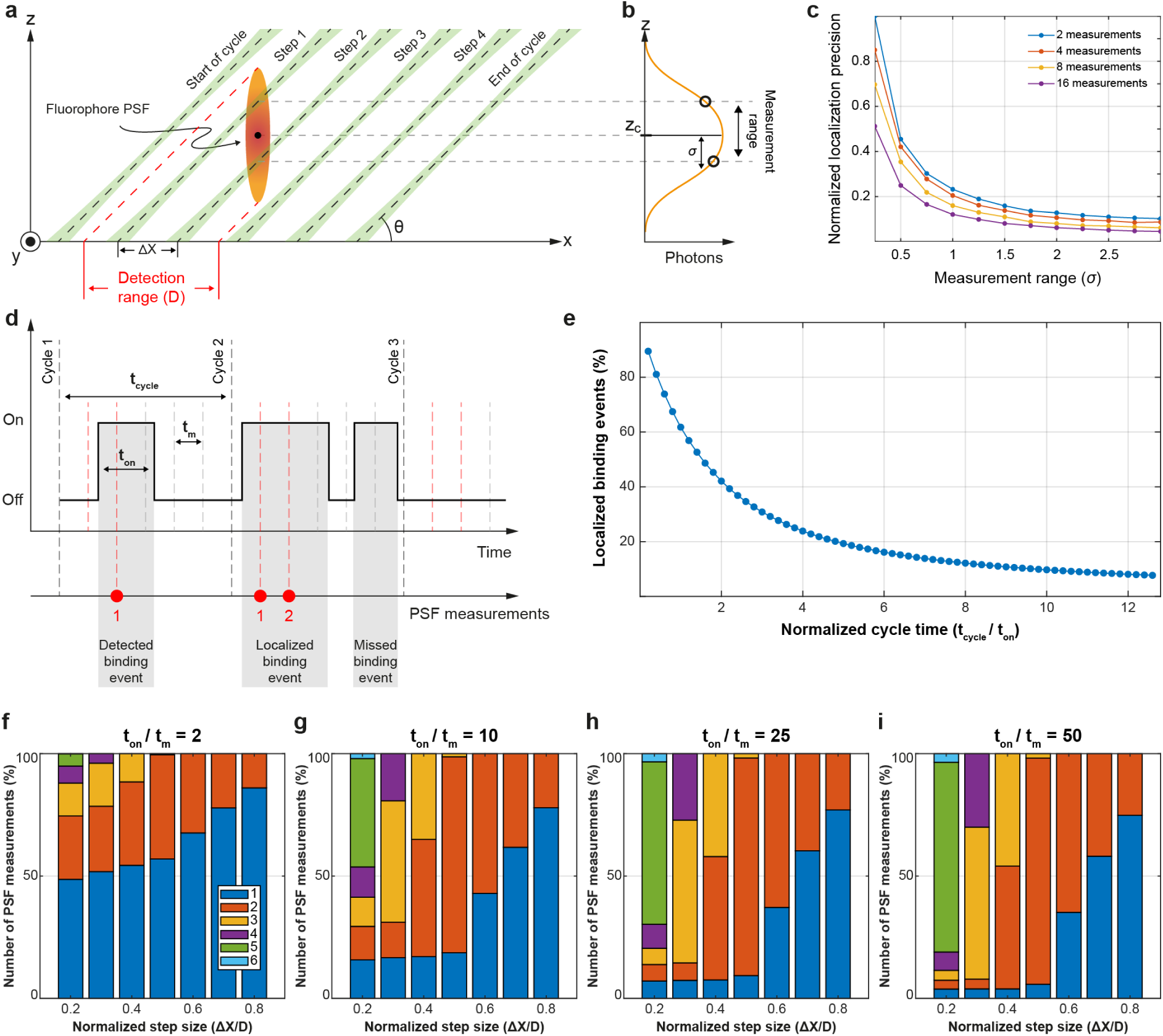
: **Simulating the effect of step-size, scan length, and exposure time. (a)** An oblique light-sheet is scanned along the x-dimension with a step size of ΔX. (**b**) As the light-sheet steps within the detection range (D) of an emitter it excites a detectable number of photons with a gaussian distribution centred at the location of the molecule. (**c**) The photon measurements from (b) can be used to localize the position of the molecule in Z and the localization precision improves with increasing measurement range and number of measurements. The measurement range is normalized to the standard deviation of the gaussian distribution shown in (b) and is measured in units of (σ). Localization precision is normalized to the maximum value. (**d**) The time course of the imaging experiment shown in (a) where imager strands transiently bind to the location of the molecule of interest with an average on time (**t_on_**) while the light-sheet is stepped along the x dimension over several imaging cycles. The vertical dashed lines indicate the time points at which the light-sheet is stepped to its next position. A red dashed line indicates that if an imager strand binds a blinking event will be produced at this time point. The dwell time of the light-sheet at each step is (**t_m_**) while the time it takes to complete one imaging cycle is (**t_c_**). Depending on the position of the light-sheet in the imaging cycle, a random binding event can be missed, produce a single measurement, or a sequence (≥2) of measurements. (**e**) The fraction of binding events that produce a sequence of ≥2 measurements and can be localized, as a function of cycle time (**t_c_**). The cycle time is normalized to the average duration of a binding event (**t_on_**). Step size: 0.5 x detection range (D), **t_on_/t_m_**=50. (**f-i**) The percentage distribution of the number of measurements collected during a binding event as a function of step size for different **t_on_/t_m_** ratios. The step size is normalized to the detection range.

**Supplementary Figure 2.**
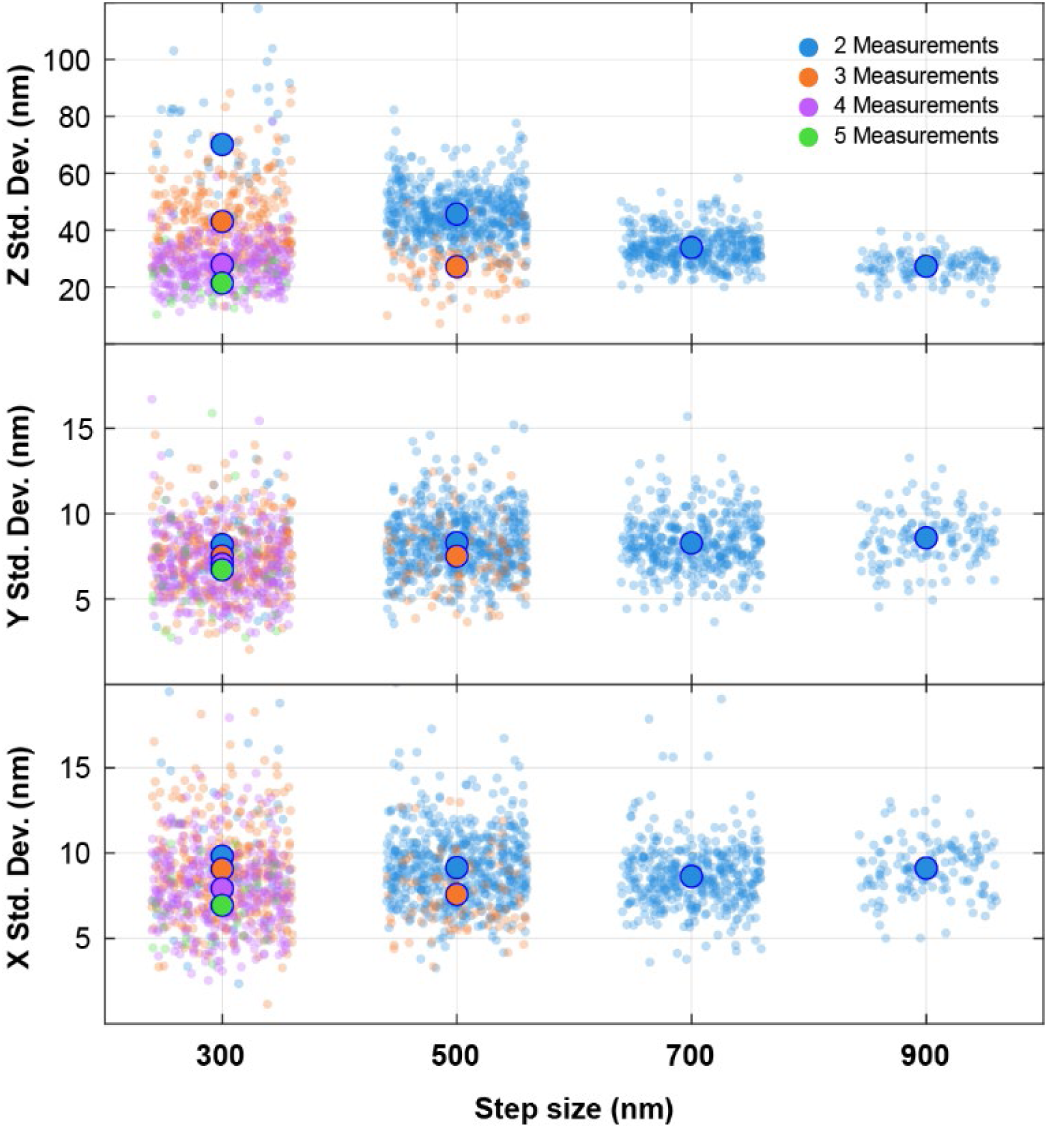
: Measuring the effect of step-size, and number of measurements on localization precision using DNA origami. X, Y and Z standard deviations as a function of step-size for different numbers of measurements. The Z standard deviation is minimized when combinations of step-size and number of measurements produce longer measurement ranges. The X and Y standard deviations appear insensitive to step size but show improvement with increasing number of measurements. Finally, we note that the X standard deviation is on average ∼**5**% worse than the Y standard deviation. This is consistent with the anisotropy in the FWHM values between the X and Y dimension of the system PSF, which is attributed to light clipping by the tertiary objective along the X dimension.

**Supplementary Figure 3.**
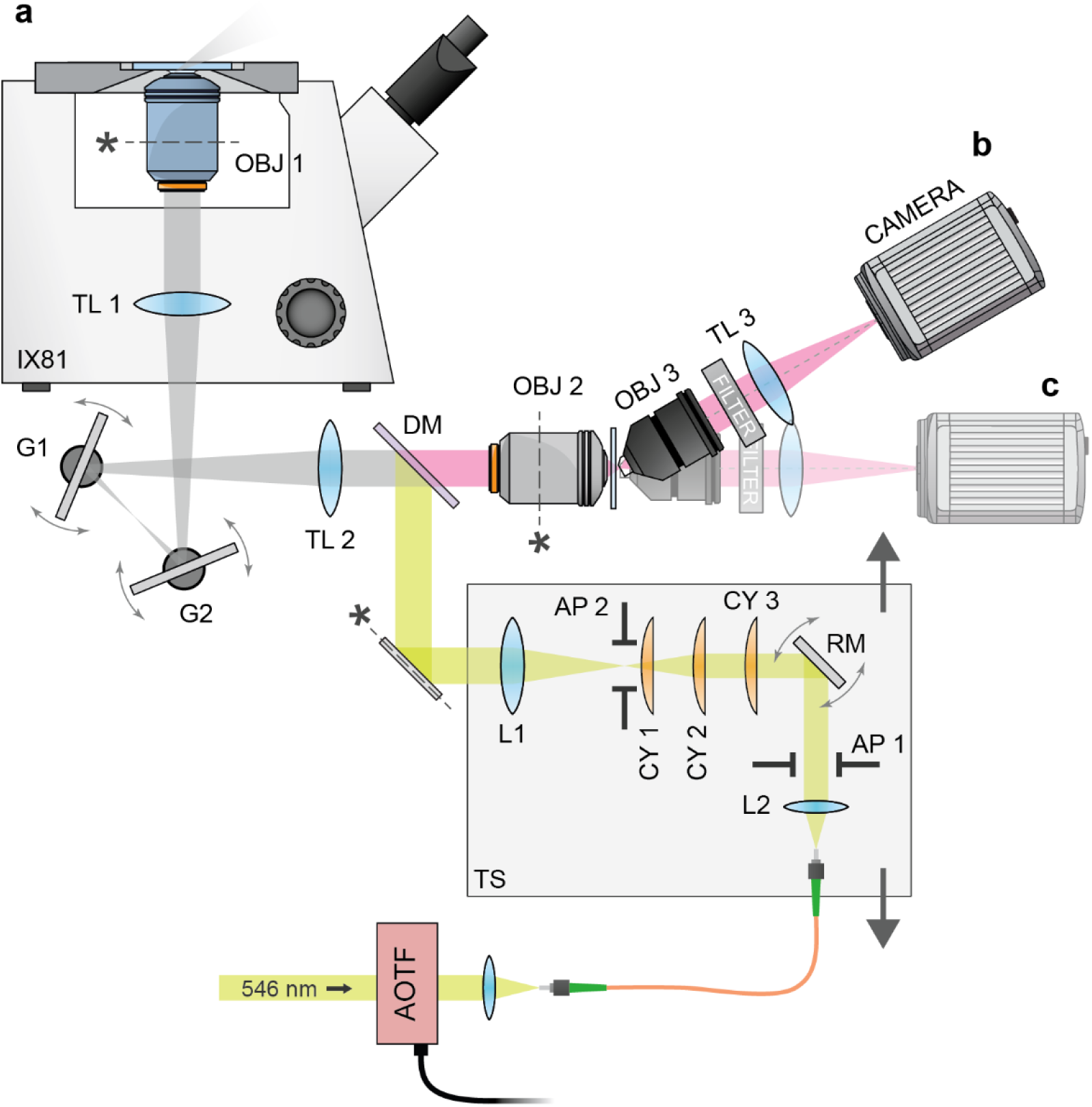
: Optical Setup. **(a)** A 546 nm CW excitation laser was passed through an AOTF and coupled into a single mode optical fibre. Light exiting the fibre was collimated **(L2)** and reflected from a resonance mirror **(RM)** for light-sheet dithering. Next, a series of cylindrical lenses were used to expand the beam size in one dimension **(CY1 and CY3)** and create a light-sheet in the orthogonal dimension **(CY2)**. Adjustable apertures controlled the light-sheet thickness (**AP 1**) and the sheet extent in the field of view (**AP 2**). Lenses **L1** and **TL2** created an image of the light-sheet at the microscope base’s (**IX81**) primary image plane. Excitation light was brought into the common light path with a dichroic mirror (**DM**). Two galvo mirrors were used for scanning the light-sheet **(G1 and G2)** while **TL1** and the primary objective **(OBJ 1)** relay the light-sheet into the sample space. The primary objective collected fluorescence from the sample volume and a remote volume was created with **TL 2** and the remote objective **(OBJ 2)**. An oblique plane in the remote volume was imaged onto the primary camera **(CAMERA)** with the combination of a tertiary objective **(OBJ 3)** and a final tube lens **(TL 3)**. The system may be used on the oblique path **(b)** as described in the text or on a straight path **(c)** when optical sectioning is not required.

**Supplementary Figure 4.**
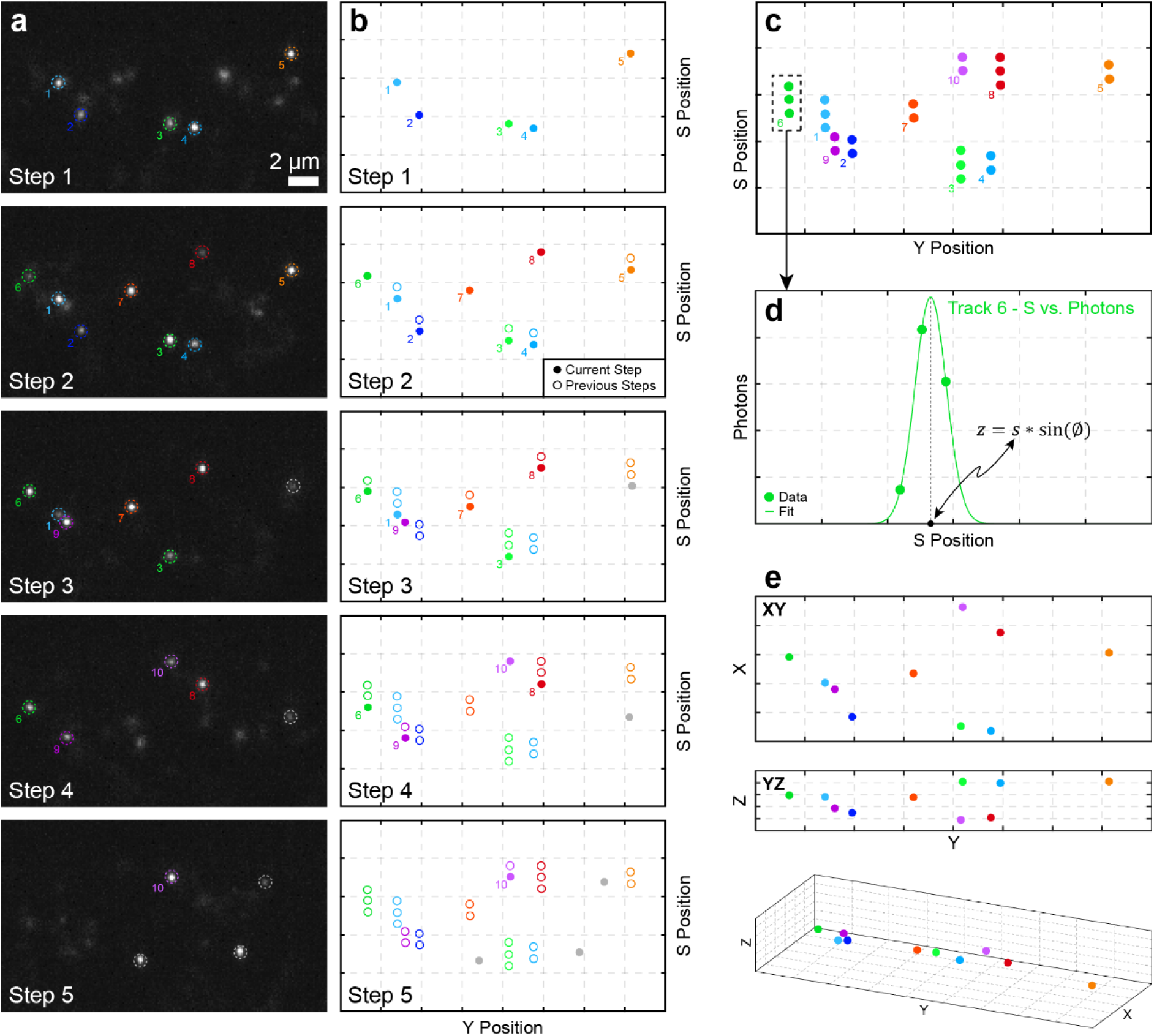
: Sequential measurements of binding events clustered into PSF tracks for (x, y, z) localization. Consecutive unprocessed camera frames are shown from top to bottom in column **(a)**. In each frame, PSF sampling locations that have been positively identified, fitted and arise from PSFs that appear in >1 frame are indicated with dashed and coloured circles and with corresponding ID numbers. PSFs only sampled once (that is, they only appear in a single frame) are shown with grey circles. Fit locations indicated on each camera frame in **(a)** are shown in an adjacent plot in column **(b)** where each plot shows the fit positions (filled circles) in the current step along with fit positions (open circles) from previous frames. As the OPM scans through the sample volume (moves forward in space and time) binding events are sampled multiple times and when plotted together as in **(c)** form “PSF tracks”. Because the distance between PSF samples is defined by the mirror step size and because OPM scanning only progresses in one direction, clustering fit positions into PSF tracks is straightforward. These PSF tracks represent multiple measurements of the same binding event sampled sequentially in space and time. The sampled s-position may be combined with the number of photons at each step to determine the z-position of a single binding event as shown in **(d)** for PSF track number 6. The (x, y) positions of each measurement within a PSF track are combined using a localization precision weighted average to determine the final (x, y) position for that binding event/PSF track as shown with XY, YZ, and 3D views in **(e)**.

**Supplementary Figure 5.**
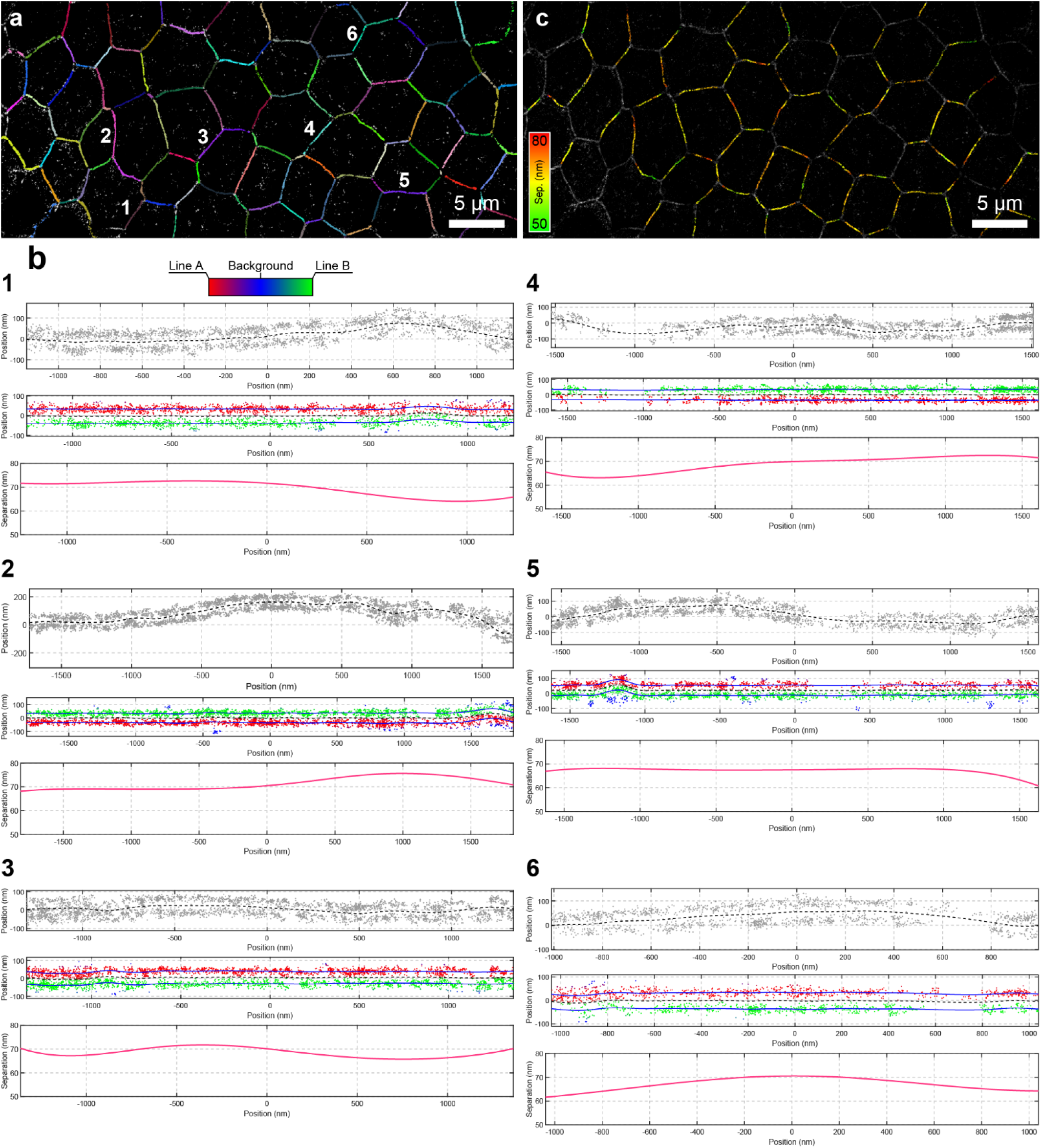
: Adherens junction segmentation and fitting workflow. **(a)** Adherens junctions (cell-cell borders) were identified from a reconstructed volume using intensity thresholding followed by watershed-based segmentation. Ridge lines derived from the watershed output were used to generate initial estimates of adherens junction positions. **(b)** Individual adherens junctions were isolated and straightened by fitting a spline to the junction centerline and transforming localization coordinates into a Frenet coordinate system. Six examples are shown from the corresponding number locations in **(a)**. Local membrane roll was then corrected to maximize separation of the opposing membrane-associated localization populations. The straightened junctions were subsequently modeled as parallel double-line structures using a Gaussian mixture model optimized by expectation-maximization (EM). The model jointly fit a central spline describing the junction centerline and a second spline describing the position-dependent separation between the opposing sides of the junction. During fitting, localizations were probabilistically assigned to one of the two junction-associated populations or to a background component. **(c)** The fitted separation values were used to generate a color-coded separation map, enabling visualization of nanoscale variations in adherens junction separation throughout the image.

